# Canonical Wnt Signaling Maintains Human Mesenchymal Progenitor Cell Multipotency During Adipose Tissue Development

**DOI:** 10.1101/2022.07.18.500487

**Authors:** Zinger Yang Loureiro, Shannon Joyce, Javier Solivan-Rivera, Anand Desai, Pantos Skritakis, Qin Yang, Tiffany DeSouza, Tammy Nguyen, Ormond A MacDougald, Silvia Corvera

**Author notes:** Corresponding author at, Program in Molecular Medicine, 373 Plantation Street, Worcester, MA 01605, USA.

## Abstract

Tissue development and repair throughout life depends on the availability of multipotent mesenchymal stem/progenitor cells capable of differentiating into specialized cell types. How an appropriately sized pool of such multipotent progenitors is maintained under varied signals for tissue growth and repair is unknown. We addressed this question by monitoring fate trajectories of human adipose tissue-derived multipotent progenitor cells using single-cell transcriptomics. Homogenous multipotent progenitors underwent two distinct fate trajectories rapidly upon induction of adipose differentiation– one toward the adipocyte fate, and the other towards a distinct, non-differentiated state characterized by up-regulation of canonical Wnt target genes. Upon isolation, this latter cell population was able to resume proliferation and display multipotency. Using canonical Wnt agonists and antagonists we find Wnt signaling is required for the maintenance of this multipotent pool under differentiation stimulus. *In vivo*, these cells are retained in adipose tissue developed from human multipotent progenitor cells in immunocompromised mice, and their transcriptomic signature is detected in human adult adipose tissue. Our study reveals a previously unrecognized mechanism for maintaining a functional pool of human mesenchymal progenitor cells under conditions of differentiation pressure, driven by Wnt signaling.

## INTRODUCTION

Adult somatic tissues contain specialized cells whose specific properties define organ- and tissue-specific functions. Replacement of these specialized cells when damaged or dead is essential for continuous tissue and organ function throughout the lifetime, and depends on the availability of multipotent stem/progenitor cells capable of differentiating into characteristic cell types. The properties of progenitor cells in epithelial tissues, such as the skin and intestine, and in blood have been well characterized ^1-3^, but how mesenchymal progenitor cells involved in the development of bone, cartilage, and adipose tissues are maintained is less clear. These mechanisms are particularly intriguing bacause a large proportion of these cells reside in adipose tissue ^4,5^, which is uniquely capable of massive expansion in adults ^6,7^. Indeed, in severe obesity, over 50% of body mass can be comprised of adipose tissue ^8^. How an adequate pool of multipotent mesenchymal progenitors is maintained under conditions of chronic differentiation pressure into the adipocyte fate is not known.

Foundational insights into mechanisms underlying adipocyte differentiation have been obtained primarily in mouse models ^9^. Two stages of murine adipocyte formation have been defined– the determination phase and the terminal differentiation phase. In the determination phase, multipotent mesenchymal progenitor cells give rise to pre-adipocytes, which remain morphologically indistinguishable from progenitors but lose the ability to differentiate into other cell types. In the terminal differentiation phase, pre-adipocytes express genes for lipid transport and synthesis, form large and specialized lipid droplets, and secrete adipocyte specific cytokines such as adiponectin. Terminal differentiation is transcriptionally controlled through sequential expression of members of the *CCAAT/enhancer binding protein* (*C/EBP*), *peroxisome proliferator-activated receptor* (*PPAR*) families, and the *adipocyte determination and differentiation factor-1/sterol response element binding protein 1c* (*ADD1/SREBP1c*), and can be inhibited *in vitro* by enforced expression of *Wnt10b*, which signals through the canonical Wnt pathway and blocks expression of *peroxisome proliferator-activated receptor-γ* (*PPARγ*) and *CCAAT/enhancer binding protein-α* (*C/EBPA)* ^10-12^,

Much less is known about the process of determination of multipotent mesenchymal progenitors into pre-adipocytes, largely due to the lack of suitable cell models. However, an important role for Wnt signaling is supported by the finding that *Wnt10b*-null mice display a progressive loss of adipogenic and osteogenic progenitors and premature adipogenesis/osteogenesis ^13^. Moreover, there is a genetic associations between mutations in *TCF7L2* and *WNT5B* and the development of type 2 diabetes ^14-16^ and between variants of *WNT10B* and the development of obesity ^17^.

Previous work has shown that mesenchymal progenitor cells in adipose tissue reside in close association with the microvasculature ^18-23^, and we have previously found that culture conditions that promote angiogenesis also promote the proliferation of mesenchymal progenitor cells ^24,25^ which give rise to multiple human adipocyte subtypes ^26^. In this study, we sought to leverage these cells to investigate the mechanisms that govern determination of mesenchymal prgenitors and their differentiation into diverse human adipocyte subtypes. Using single cell transcriptomics, we find that upon adipogenic stimulation, human multipotent mesenchymal progenitor cells differentiate into adipocytes, but there is a simultaneous induction of a pool of cells that do not differentiate. Upon isolation, these cells regain proliferative capacity and the ability to differentiate into multiple lineages, including chondro- and osteo-genic lineages. This multipotent reservoir is characterized by expression of Wnt target genes, and its size is controlled by canonical Wnt signaling. These results reveal a mechanism, elicited under conditions of strong differentiation pressure, that maintains a pool of functional multipotent mesenchymal progenitors, and explains how human mesenchymal tissues can be maintained and repaired throughout the lifetime.

## RESULTS

### Acute transcriptional remodeling of multipotent progenitor cells in response to adipogenic stimulation

Small fragments of human subcutaneous adipose tissue from subjects undergoing elective panniculectomy surgery were harvested within six hours of surgery and embedded in Matrigel. After 14 days in culture, extensive growth of capillary sprouts was observed (Fig. 1a, top panel). Sprouts were digested using Dispase and Collagenase Type I and plated in plastic culture dishes, where they adopted a fibroblastic homogenous phenotype characteristic of mesenchymal progenitor cells (Fig. 1a, right panel). After two passages, cells were frozen for further studies. To determine whether cells obtained by this method retain multipotency, monolayers were exposed to adipose, chondro- or osteo-genic differentiation media for 10 day and stained or subjected to bulk RNA sequencing. Staining for neutral lipid, proteoglycan or calcium was seen only in cells exposed to specific differentiation cocktails (Fig. 1b). Evidence for multipotent differentiation was also seen in gene expression profiles, where selected genes associated with the adipogenic (*ADIPOQ, PLIN1*), chondrogenic (*ACAN, COL10A1*) or osteogenic (*ALPL, SMOC2*) lineages were selectively expressed in response to each cocktail (Fig. 1c). These results confirm that mesenchymal progenitor cells derived by this method are multipotent. Notably, not all cells underwent differentiation, as cells lacking lipid droplets could be detected alongside lipid replete cells (Fig. 1b, top panel).

**Figure. 1.**
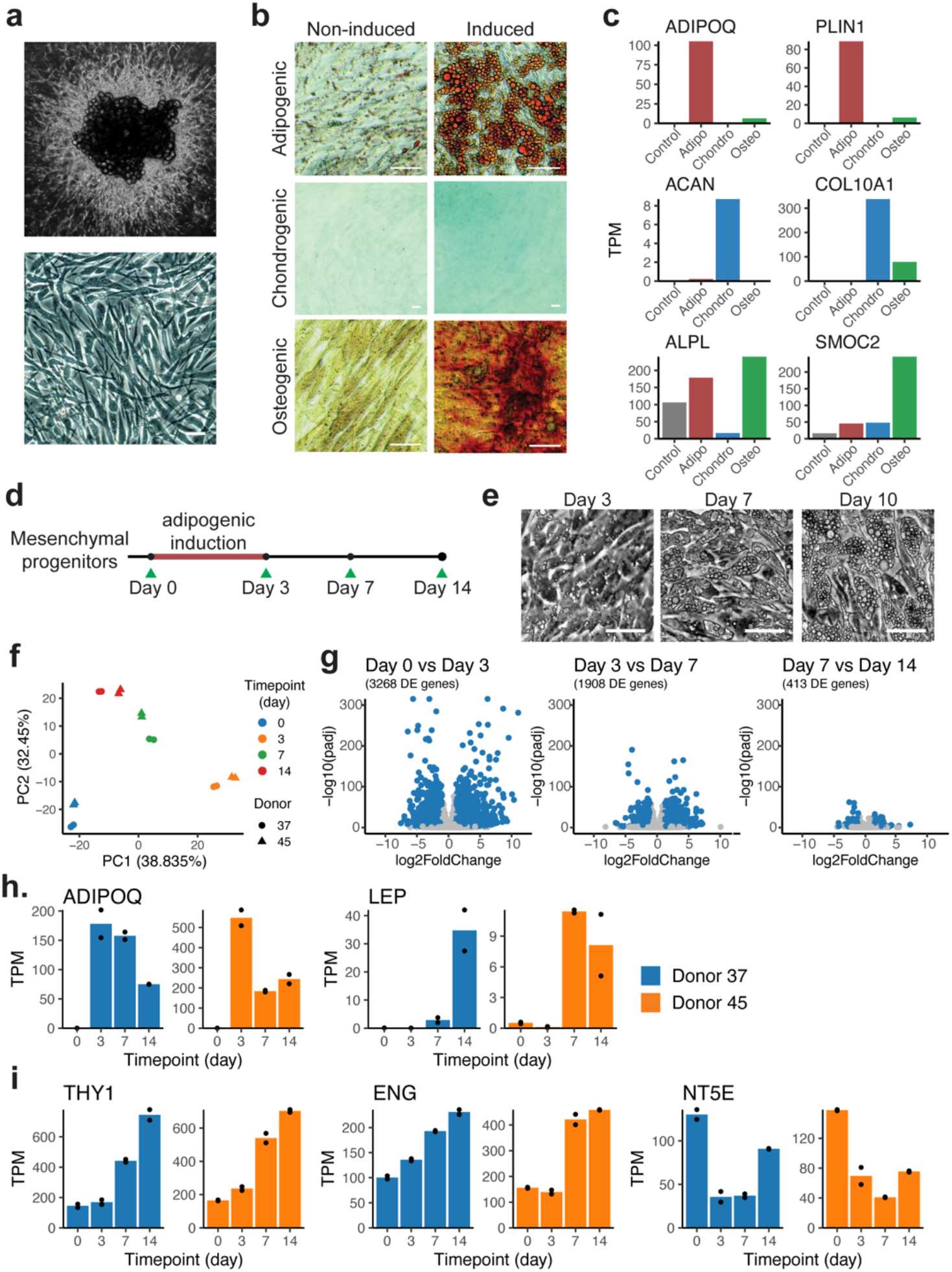
Dynamic transcriptomic changes in multipotent mesenchymal progenitors from human adipose tissue undergoing adipogenic differentiation. **a**. Mesenchymal progenitor cells expanded from adipose tissue explants in 3-dimensional culture (top panel), plated grown to confluence in 2-dimensional culture dishes (bottom panel; scale bar, 50 µm). **b**. Images of progenitors induced toward adipogenic, chondrogenic or osteogenic cell fates for 10 days. Adipogenic-induced cells were stained with Oil Red O, chondrogenic-induced cells with Alcian Blue 8GX, and osteogenic-induced cells with Alizarin Red S. Scale bar, 50 µm. **c**. Marker genes for progenitors differentiated toward adipogenic, chondrogenic, and osteogenic lineages identified using their transcriptomic profile. Bars are means of technical replicates from n=1 wells subjected to the indicated differentiation cocktails. *ADIPOQ: Adiponectin, PLIN1: Perilipin 1, ACAN: Aggrecan, COL10A1: Collagen Type X Alpha 1 Chain, Chondrocyte specific, ALPL: Alkaline Phosphatase, SMOC2: SPARC/Osteonectin-Related Modular Calcium-Binding Protein 2*. **d**. Schematic of the adipogenesis time-course study. **e**. Representative images of mesenchymal cells induced toward adipogenic fate for 3, 7, and 10 days. Scale bar, 50 µm. **f**. Scatter plot of the first two principal components of bulk RNASeq results from two independent cultures, each from two independent donors, separately expanded, and used to obtain RNA at 0, 3, 7, or 14 days post adipogenic induction. Principal component analysis (PCA) was performed on the expression of the top 1000 most variable genes across all n=16 samples. **g**. Volcano plots of the differential gene expression analysis results between consecutive time points. **h**,**i**. Time courses of adipokines *Adiponectin (ADIPOQ)* and *Leptin (LEP)*,and of mesenchymal progenitor markers *THY1 (CD90), ENG (CD105)*, and *NT5E (CD73)*.

To further understand mechanisms governing adipogenic fate commitment, we performed bulk RNA-seq to analyze the transcriptomes of progenitors derived from two independent donors at 0, 3, 7 and 14 days post adipogenic induction (Fig. 1d). Principal component analysis (PCA) (Fig. 1f) reveals the largest variance to be associated with differentiation and little variance attributable to the tissue donor. The largest gene expression changes occur between 0 and 3 days of exposure to adipogenic induction, when the cells have not yet accumulated large lipid droplets (Fig. 1e). Consistently, the number of differentially expressed genes between 0 and 3 days of differentiation (3268 genes) is higher than that seen between 3 and 7 days (1908 genes), or between 7 and 14 days (413 genes) (Fig. 1g). Expression of adiponectin (*ADIPOQ*), an adipocyte specific cytokine, is maximal at 3 days after induction (Fig. 1h), indicating that commitment to the adipogenic fate occurs early. Additional adipocyte development continues beyond 3 days, as evidenced by delayed expression of leptin (*LEP*), another adipocyte-associated cytokine (Fig. 1h). Concomitantly with induction of adipocyte genes, mesenchymal progenitor cell markers *THY1/CD90, ENG/CD105, NT5E/CD73* were detected at all time points, and *THY1* and *ENG* expression increased over time (Fig. 1i). The detection of mesenchymal progenitor and differentiated adipocyte markers is consistent with a heterogenous cell population during the differentiation process.

### Single-cell RNA-seq reveals adipogenic induction elicits two distinct fates trajectories

To understand the specific transcriptomic changes in cells undergoing adipogenic differentiation without confounding signals from cells that do not undergo differentiation, we performed single cell RNA-seq on two separate cultures: one corresponding to multipotent progenitors grown to confluency but not subjected to differentiation stimuli, and the second corresponding to progenitors exposed to adipogenic media for 3 days (Fig. 2a, Extended Data Fig. 1). At this time point, cells displayed minimal lipid accumulation and were thereby still amenable to the microfluidic-based single cell profiling. Projection of cells from these two culture conditions by the top two principal components showed the two populations were non-overlapping (Fig. 2b), indicating that all cells undergo extensive transcriptomic changes upon adipogenic induction. Interestingly, a broader transcriptomic spectrum is seen in cells subjected to adipogenic induction, as evidenced by the larger spread in the principal component projection.

**Figure 2.**
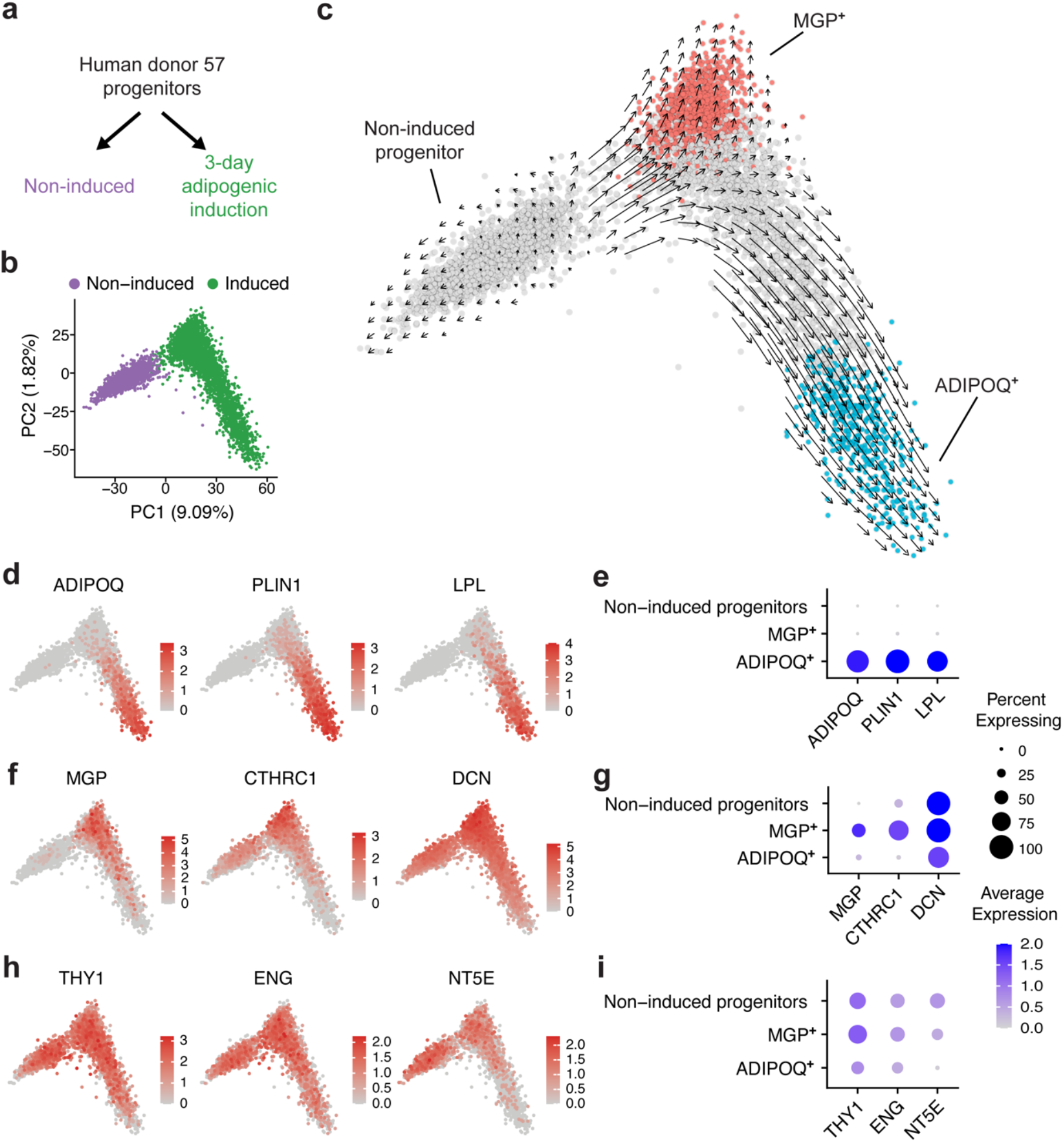
Single-cell RNA-seq of induced adipose progenitors. **a**. Schematic of the early adipogenesis single-cell transcriptomic profiling study. **b**. PCA projection of the single-cell profile of 6615 cells (3226 non-induced, 3430 adipogenic-induced, mean number of genes per cell = 3382). **c**. Inference of developmental trajectory with RNA velocity. Red and blue colored cells represent clusters at the terminals of two projected fate trajectories. **d**,**e**. Representative adipocyte marker genes in PCA projection and dot plots. *ADIPOQ: Adiponectin, PLIN1: Perilipin, LPL: Lipoprotein Lipase*. **f**,**g**. Representative *MGP*^+^ cluster marker genes in PCA projections and dot plots. *MGP: Matrix Gla Protein, CTHRC1: Collagen Triple Helix Repeat Containing 1, DCN: Decorin*. **h**,**i**. Mesenchymal progenitor marker genes in PCA projections and dot plots.

To gain insight on the nature of the transcriptomic variance, we performed developmental trajectory inference using RNA velocity. This analysis indicates that upon induction, progenitors diverge along distinct trajectories toward two cell fates (Fig. 2c): Cells in the terminal of one trajectory expressed adipocyte-specific genes including *adiponectin* (*ADIPOQ*), *perilipin* (*PLIN1*), and *lipoprotein lipase* (*LPL*), while cells at the terminal of the opposite trajectory expressed extracellular matrix genes (*MGP/matrix gla protein, DCN/decorin, CTHRC1/collagen triple helix repeat containing 1*) (Fig. 2d-g). Importantly, these cells also retained mesenchymal progenitor marker expression (*THY1, NT5E, ENG*) (Fig. 2h,i). The observation of two fates upon adipogenic induction was consistent with morphological observations that, upon adipogenic induction, a fraction of cells routinely failed to accumulate lipid droplet (Fig. 1b). To evaluate whether the markers identified were specific to adipocyte differentiation or preserved across different lineages, we profiled the transcriptome of progenitors induced toward adipogenic, chondrogenic, or osteogenic lineages for 3 days (Extended Data Fig. 2a-d, 3c-d).

Adipocyte-specific genes *ADIPOQ, PLIN1, LPL* were consistently detected in the adipogenic induced cells in all profiled datasets (Extended Data Fig. 3a,c), but *CTHRC1* and *DCN*, identified in cells of the other trajectory, were detectable in non-induced cells in other profiled datasets (Extended Data Fig. 3b,d), suggesting that these genes may not be representative fate markers. We selected *MGP* as a representative marker for the induced, non-adipogenic population, as *MGP* expression was not detected in uninduced progenitors and was consistently upregulated in adipogenic induction cells from all donors analyzed (Extended Data Fig. 3b,d). Intriguingly, in the transcriptomic profile of progenitor cells stimulated with adipogenic/chondrogenic/osteogenic induction or basal media for three days, *MGP* was upregulated in all induced conditions (Extended Data Fig. 3d), suggesting the *MGP*^+^ cells may represent a generic population that is intentionally preserved.

### MGP^+^ cells express Wnt target genes, retain proliferative potential and multipotency

To further investigate the identity of the *MGP* ^*+*^ cells, we conducted differential gene expression analysis comparing the *MGP* ^*+*^ and *ADIPOQ*^*+*^ cells (Fig. 3a). As expected, *ADIPOQ*^*+*^ cells were enriched in genes associated with adipocyte-related pathways (Fig. 3b). In contrast, *MGP* ^*+*^ cells were enriched in genes associated with skeletal development pathways, mostly comprising extracellular matrix proteins (Fig. 3c). However, these cells did not express osteocyte (*osteocalcin*/*BGLAP, osteopontin*/*SPP1*) or chondrocyte (*aggrecan*/*ACAN, cartilage collagen*/*COL2A1*) markers, suggesting that the *MGP*^+^ cells were not comprised of cells undergoing differentiation into an alternative mesenchymal cell lineage. Further analysis of the *MGP*^+^ population revealed that multiple canonical Wnt target genes (*SFRP2, DPP4, DKK1, SNAI2, WISP2*) were significantly up-regulated (Fig. 3d,e). Components of canonical Wnt signaling including Wnt ligands *WNT5A, WNT5B* and core pathway members *β-catenin/CTNNB1, TCF7L2* were expressed (Fig. 3f), suggesting the cells were capable of Wnt activities. The RNA-seq results of primary mouse mesenchymal progenitors with *β-Catenin* knockout or *Wnt3a* treatment from a study previously published ^10^ revealed *Mgp* level decreased in *β-Catenin* knockout cells comparing to the control and *Mgp* level increased after *Wnt3a* treatment in both tested timepoint (Fig.3g), implying *MGP* is a downstream gene of canonical Wnt signaling. Among the Wnt target genes, *DPP4* was reportedly a marker for undifferentiated adipocyte progenitor ^27^. Treatment of progenitors with *DPP4*-specific inhibitors LAF237 or MK0431 during adipogenic induction do not increase adipocyte lipid droplet number or sizes (Extended Data Fig. 4), indicating *DPP4* did not have a functional role in progenitor maintenance. Moreover, ligand-receptor analysis of potential interactions between *ADIPOQ*^+^ and *MGP*^+^ cells revealed Wnt-induced *Ephrin B1* (*EFNB1*) in *ADIPOQ*^*+*^ cells and *Ephrin type-B receptor 6* (*EPHB6*) in the *MGP*^*+*^ cells (Fig. 3h,i). Other significant interactions between *ADIPOQ*^*+*^ and *MGP* ^*+*^cells included of collagens, integrins and other structural extracellular proteins interacting with cognate receptors (Supplemental Table 1). These results indicate that *MGP*^*+*^ cells comprise a unique population of cells with no clear mesenchymal lineage fate commitment, and they were responsive to Wnt signal.

**Figure 3.**
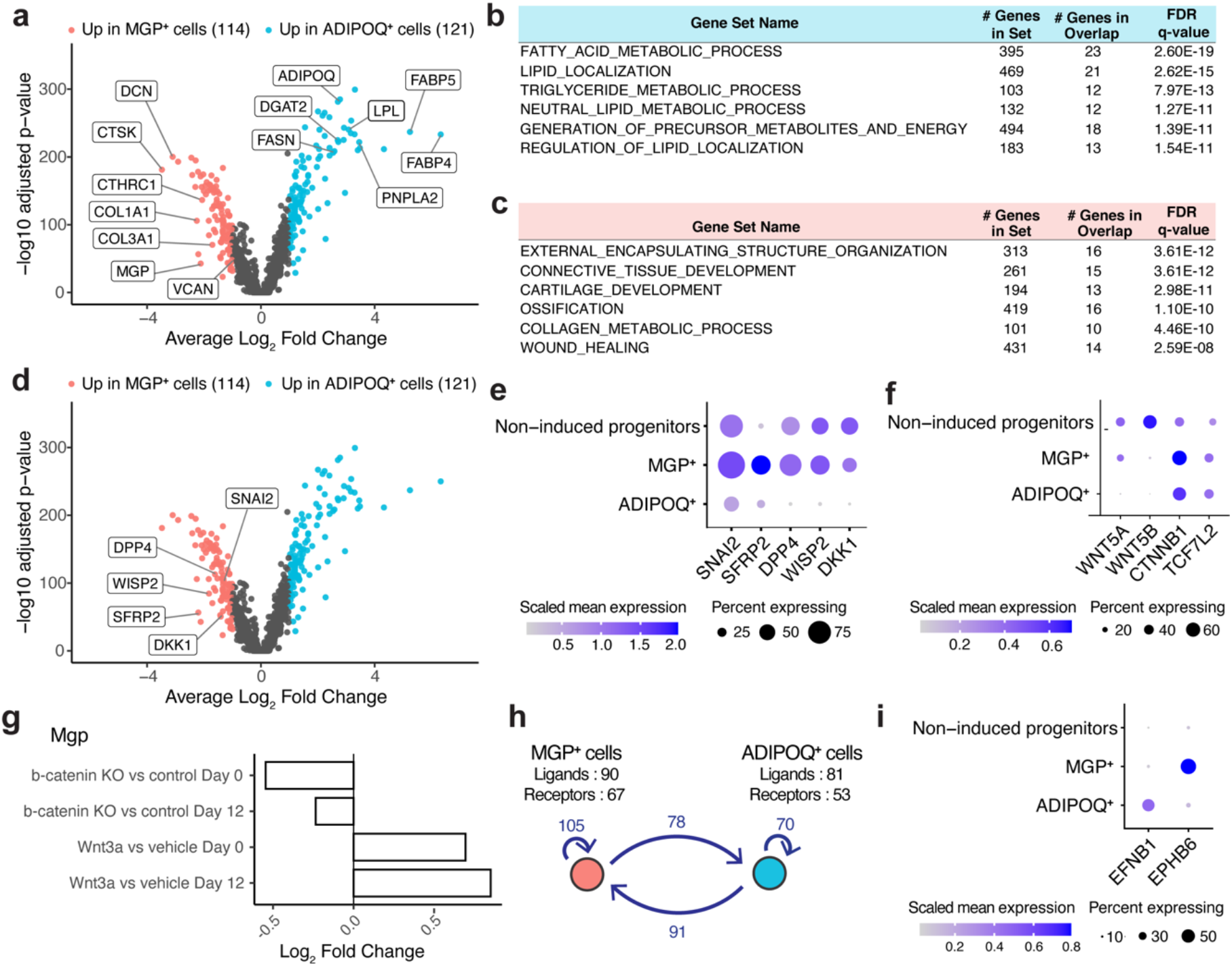
Active canonical Wnt signaling in *MGP*^+^ cells . **a**. Volcano plot comparing *MGP*^+^ and *ADIPOQ*^+^ cell gene expression, with most differentially expressed genes highlighted. Tested features were limited to genes detected in >25% cells in at least one of the clusters. Differentially expressed genes were defined as those with log2 fold change > 1 and adjusted p-value < 0.001. **b**. Top 6 significantly enriched gene sets of genes up-regulated in the *ADIPOQ*^+^ cells. **c**. Top 6 significantly enriched gene sets of genes up-regulated in the *MGP*^+^ cells. **d**. Identical volcano plot as **a**, with canonical Wnt target genes highlighted. **e**. Dot plot of the canonical Wnt target genes that were significantly up-regulated in *MGP*^+^ cells. **f**. Dot plot of the Wnt ligand and core Wnt pathway members. **g**. bar graph of log_2_ fold change values of *Mgp* from four separate differential expression analyses of primary mouse mesenchymal progenitors isolated directly from *β-Catenin* ^fl/fl^ mice. Non-adipogenic-induced progenitor (annotated as “Day 0”) or progenitors underwent 12-day adipogenic induction (annotated as “Day 12”) were harvested for RNA-seq under Wnt perturbations: Day 0 or Day 12 cells were either induced for *β-Catenin* knockout (annotated as “KO”) or were treated with 20 ng/ml recombinant Wnt3a or vehicle for 4 hours before harvest (*n*=4 per group). **h**. Ligand-receptor analysis identifies multiple ligand-receptor interacting pairs between the *MGP*^+^ and *ADIPOQ*^+^ cells. **i**. Dot plot of *Ephrin-B Receptor 6* (*EPHB6*) and *Ephrin B1* (*EFNB1*) in *MGP*^+^ or *ADIPOQ*^+^ cells.

To further investigate the functional characteristics of the *MGP* ^*+*^cells, we first tested whether they represent the cells that fail to accumulate lipid droplets after exposure to adipogenic induction signal. The presence of lipid droplets decreases cell density, and 7-day adipogenic induced cells could be separated by centrifugation through Percoll gradients (Fig. 4a). Cells from cultures that were not exposed to differentiation induction were recovered between 1.02-1.04 g/ml densities (referred to as high-density cells), while cells differentiated for 7 days were recovered in two populations, one at the 1.01-1.02 g/ml (low-density cells) and another between 1.02-1.04 g/ml densities (high-density cells). The induced cells in the low-density population contained visible lipid droplets, while the majority cells in the high-density population did not (Fig. 4a). To determine whether cells in the high-density population represented *MGP*^+^ cells, we conducted bulk RNA-Seq of the high- and low-density induced cells and reviewed the expression of *ADIPOQ*^+^ and *MGP*^+^ cell markers, as determined by the single-cell RNA-Seq. We found that *ADIPOQ*^+^ cells were highly enriched in the low-density cells, while *MGP*^+^ cells were enriched in the high-density population (Fig. 4b). qRT-PCR confirmed a strong enrichment of *ADIPOQ* and *MGP* in induced, low- and high-density cells, respectively (Fig. 4c).

**Figure 4.**
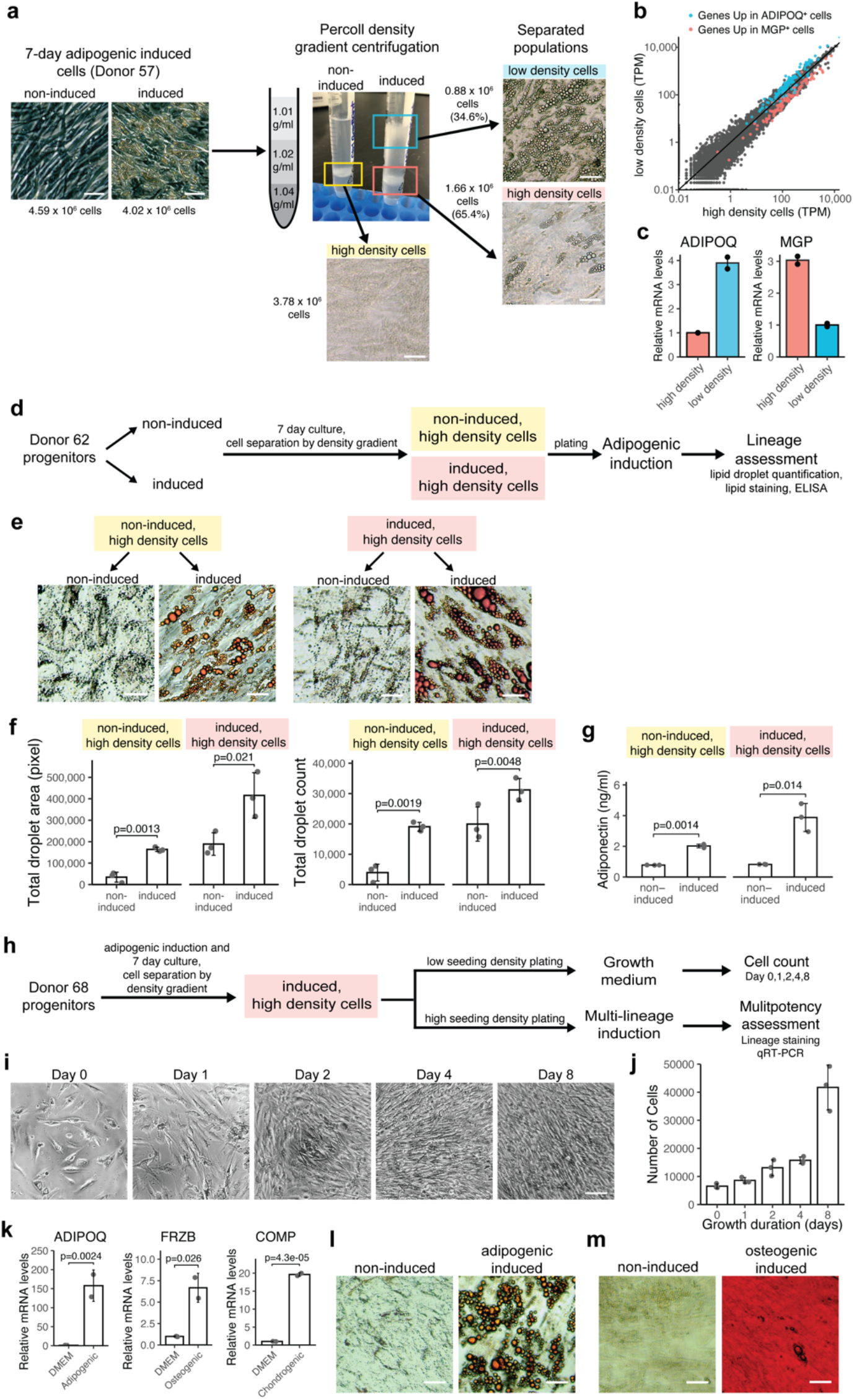
*MGP*^+^ cells are multipotent mesenchymal progenitors. **a**. Schematic and images of the cell separation assay. Non-induced or 7-day adipogenic induced cells from Donor 57 were subjected to Percoll density gradient centrifugation. Scale bars, 50 µm. **b**. Scatter plot of each detected gene’s transcript per million (TPM) values between the high- and low-density cells measured by bulk RNA-sequencing. Genes highlighted in blue were the 121 genes enriched in the *ADIPOQ*^+^ cells, as identified by the differential expression analysis of single-cell RNA-seq presented in Fig. 3a, and genes highlighted in red were the 114 genes enriched in the *MGP*^+^ cells. **c**. qRT-PCR assessment of *ADIPOQ* and *MGP* mRNA levels in the high- and low-density cells extracted from 7-day adipogenic induced cells. Plotted are means of n=2 independent experiments each assayed with technical duplication. **d**. Schematic of the experiment assessing adipogenic potential of the high density cells obtained from a density gradient centrifugation. **e**. Oil Red O staining of the high density cells after additional 7-day adipogenic induction. Scale bar 50 µm. **f**. Lipid droplet count and droplet area quantification of the high density cells after additional 7-day adipogenic induction (error bars = SD, *n* = 3, exact p-values shown were determined by unpaired two-tailed *t*-tests). **g**. Adiponectin level in the conditioned media of high density cells after additional 7-day adipogenic induction, measured by ELISA (error bars = SD, *n* = 3, exact p-values shown were determined by unpaired two-tailed *t*-tests). **h**. Schematic of the experiment assessing multipotent and proliferative potential of the high density cells obtained from a density gradient centrifugation of 7-day adipogenic induced cells. For low seeding density plating, 10000 cells were plated per well in a 96-well plate; for high seeding density plating, 30000 cells were plated per well in a 96-well plate. **i**, Phase images of the induced, high density cells after indicated days in progenitor growth medium. Scale bar, 100 µm. **j**. Cell counts of the induced, high density cells after indicated days in progenitor growth medium (error bars = SD, *n* = 3). **k**. mRNA levels of adipogenic (ADIPOQ), chondrogenic (COMP), and osteogenic (FRZB) lineage markers of induced, high density cells after 10-day lineage differentiation (error bars = SD, *n* = 2, exact p-values shown were determined by unpaired two-tailed *t*-tests). **l**. Oil Red O staining of induced, high density cells after 10-day adipogenic induction. Scale bar, 50 µm. **m**. Alizarin Red S staining of induced, high density cells after 10-day osteogenic induction. Scale bar, 50 µm.

The ability to enrich for *MGP*^+^ cells by density centrifugation allowed us to analyze their functional properties. We first queried whether these progenitor cells were inherently unable to undergo adipogenic differentiation. We grew multipotent progenitors to confluence and left one group untreated (referred to as “non-induced”) and subjected a second group to adipogenic differentiation for 7 days (referred to as “induced”). Cells were then lifted and subjected to percoll density separation. The high-density cells recovered from both the non-induced and induced populations were then seeded at high seeding density, such that confluency was achieved upon adherence (Fig. 4d). 48 hours post cell seeding, the high-density cells were subjected to additional adipogenic differentiation for 7 days. Oil red O staining and lipid droplet quantification revealed lipid droplet accumulation (Fig. 4e,f) and *ADIPOQ* was secreted (Fig. 4g) in the high-density cells recovered from either the non-induced or the induced cultures. The results revealed that *MGP*^+^, lipid-devoid cells generated upon adipogenic stimulation were responsive to adipogenic stimulus, as they were able to undergo adipogenesis upon a second round of induction.

The results above suggested that *MGP*^+^ cells might represent a reservoir of multipotent progenitors generated in response to adipogenic pressure. To test this hypothesis, we analyzed whether *MGP* ^*+*^cells retain proliferative capacity and multipotency. High density cells recovered after 7 days of adipogenic induction were plated at low seeding density in media used to expand multipotent progenitors (Fig. 4h). Within 24 hours, cells began to proliferate and maintained the spindle-like fibroblastic morphology characteristic of progenitor cells (Fig 4i,j). Additional high-density cells were exposed to either adipogenic or osteogenic differentiation induction media and analyzed by qRT-PCR after 3 days and by staining after 14 days (Fig. 4h). Lineage markers for adipo-, chondro-, and osteo-genic fate, as defined by early multi-lineage transcriptome profiling (Extended Data Fig. 2e,f), were expressed with each respective differentiation induction (Fig. 4k). Lipid droplets and calcium aggregates were detected upon adipogenic or osteogenic differentiation, respectively (Fig. 4l,m). These results indicate that a specific pool of progenitor cells retain proliferative capacity and multipotency for a minimum of 7 days after induction of adipogenic differentiation.

### The *MGP*^*+*^ cells are maintained *in vivo*

We next sought to determine whether *MGP*^+^ cells are present *in vivo* during human adipose tissue development. We leveraged a hybrid *in vivo* model, in which human mesenchymal progenitor cells were implanted into immuno-compromised mice after adipogenic induction (Fig. 5a, Extended Data Fig. 5). These cells generated a functional human/mouse hybrid adipose depot (Fig. 5b)^28^. Analysis of human transcript-specific reads in the implants 8 weeks after implantation revealed expected expression of human adipocyte genes (*ADIPOQ, PLIN1, LPL*), but also revealed expression of *MGP*^*+*^ cluster markers (*MGP, CTHRC1, DCN*) (Fig. 5c,d), suggesting that a pool of multipotent progenitors was actively maintained in an *in vivo* environment. We then selected the top 40 differentially expressed genes between the *MGP*^+^ and the *ADIPOQ*^+^ single-cell populations to generate a signature for each cell type (Fig. 5e). To determine whether *MGP*^+^ and the *ADIPOQ*^+^ cell populations were present in human adult adipose tissue, we leveraged an atlas of single-cell and single-nuclei transcriptomes recently provided by Emont, et al.^29^. Emont et al identified multiple cell types comprising adult adipose tissue, including a population of stem/progenitor cells termed ASPCs. We found a clear signature of *MGP*^+^ cells in ASPCs, and of *ADIPOQ*^+^ cells in mature adipocytes (Fig. 5f), indicating that *MGP*^+^ cells were maintained throughout development and in adult human adipose tissue. *MGP* was expressed both in ASPCs of subcutaneous (SAT) and visceral adipose tissue (VAT), with SAT having overall higher level of *MGP*^*+*^ cell marker genes (Extended Data Fig. 6a,b), consistent with SAT having higher expandability. Moreover, expression of Wnt target genes characteristic of *MGP*^+^ cells was also seen in ASPCs and was negligible in mature adipocytes (Extended Data Fig. 6c). These results indicate that the signature of *MGP*^+^ and *ADIPOQ*^*+*^ cells is stable during adipose tissue development and maintenance. To further test this we analyzed the abundance of signature genes over 14 days of adipogenic differentiation *in vitro* (Fig. 5g). We find that these genes are stable over time and remain highly expressed in cells during *in vivo* adipose tissue development (Fig. 5h).

**Figure 5.**
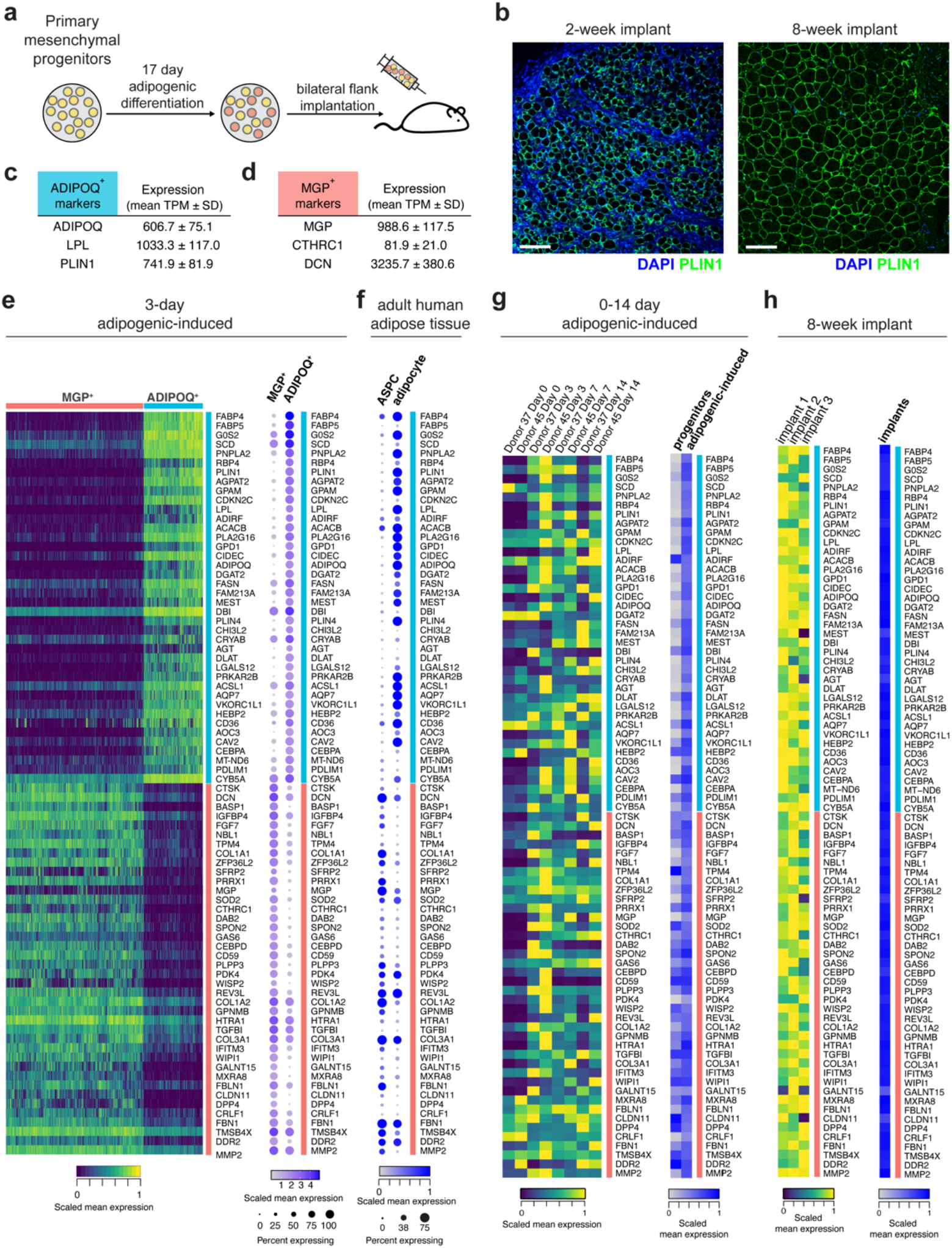
*MGP*^+^ cells are maintained over time in culture, and *in vivo*. **a**. Schematic of the human adipogenic-induced progenitor mouse implantation model. **b**. Histological sections of implants 2 weeks (left panel) and 8 weeks (right panel) after injection. **c**,**d**. mRNA levels of the *ADIPOQ*^+^ and MGP+ cell markers in the implants measured by bulk RNA-seq. **e**. Heatmap of individual cells and summary dot plot of gene expressions of the top 40 differentially expressed gene from *MGP*^+^ and *ADIPOQ*^+^ cells, derived from the single-cell dataset described in Fig.2. **f**. Dot plot of gene expression of the top 40 marker gene of *MGP*^+^ and *ADIPOQ*^+^ cells in the published single-cell/single-nuclei adipose tissue transcriptome of adult human ages 29-73 years published by Emont, et al. ^29^. **g**. Heatmap of gene expression of the top 40 differentially expressed gene of *MGP*^+^ and *ADIPOQ*^+^ cells from the adipogenesis time course bulk RNA-seq dataset described in Fig. 1d, presented as individual samples (left panel) or summarized by adipogenic induction status (right panel). **h** Heatmap of expression of the top 40 differentially expressed gene of *MGP*^+^ and *ADIPOQ*^+^ cells from the human reads of the implant dataset presented in Fig. 5a, presented as individual samples (left panel) or summarized (right panel).

### Wnt signaling controls homeostasis of *MGP*^+^ and *ADIPOQ*^+^ cell pools

The expression of Wnt target genes in *MGP*^+^ cells and ASPCs raises the possibility that Wnt signaling plays an active role in the maintenance of multipotent progenitor cells. To test this possibility, we examined the effects of canonical Wnt signaling during the early stages of adipogenic induction. We first analyzed the effects of the canonical Wnt agonist CHIR99021, which acts by inhibiting *GSK3* and preventing phosphorylation-triggered degradation of *β-Catenin*. In preliminary experiments, we found that chronic treatment of cells undergoing differentiation with CHIR99021 resulted in a decrease in ATP levels detectable at concentrations above 0.5 uM and increase in membrane permeability reflective of compromised cell viability (Fig. 6a, Extended Data Fig. 7). Accordingly, we used a low dose (0.4 uM) to examine the effects of Wnt activation in progenitor cells on the extent of subsequent adipogenic differentiation, assessed by measuring the number and size of lipid droplets at day 9 after induction (Fig. 6b) as well as expression of *ADIPOQ* at day 3 after induction. One day of exposure to CHIR99021, before addition of the adipogenic cocktail, resulted in a significant decrease in the number and size of lipid droplets at day 9 (Fig. 6c,d). Expression of *ADIPOQ* was progressively suppressed with 2 and 3 days of exposure (Fig. 6e). To test whether suppression of canonical Wnt signaling would have the converse effect, we used the same paradigm to test the effects of XAV939, which stabilizes *AXIN* and promotes destruction of *β-Catenin*. An increase in lipid droplet number and size (Fig. 6g,h) and an increase in *ADIPOQ* levels (Fig. 6i) was seen in response to 1-2 days of exposure to the Wnt signaling inhibitor. Notably, CHIR990021 or XAV939 treatment resulted in *MGP* expression level changes in opposite direction to expression level changes of *ADIPOQ* (Fig 6f,j). These results suggest that activation of canonical Wnt signaling shifts the cell fate preference towards maintenance of an undifferentiated progenitor pool.

**Figure 6.**
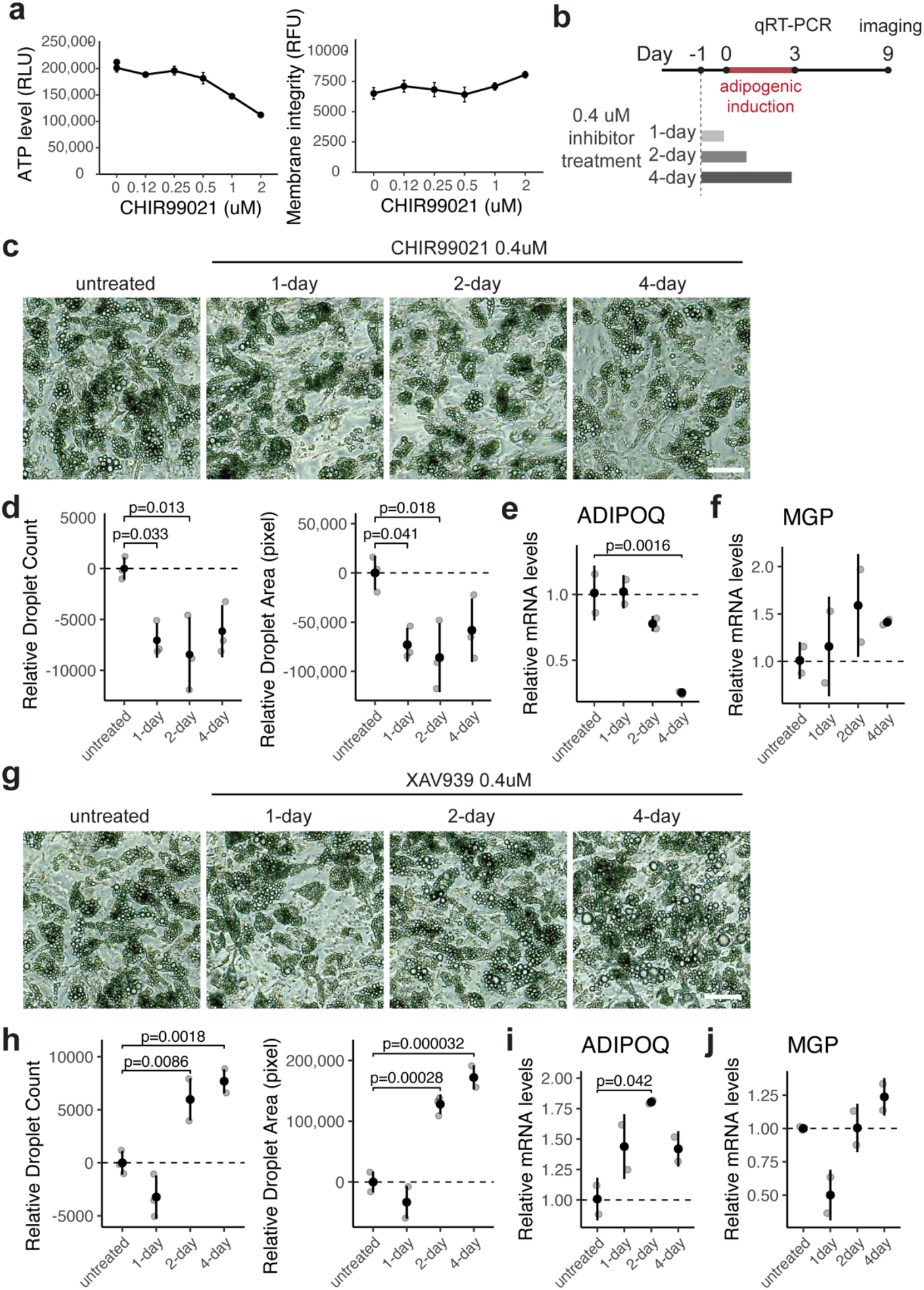
Canonical Wnt signaling controls cell fate balance between adipogenic differentiation and progenitor maintenance. **a**. ATP levels (left panel) and membrane permeability (right panel) in 10-day adipogenic-induced cells exposed every 24h to different CHIR99021 at the concentrations indicated (error bars = SD, *n =* 3). **b**. Schematic of the assay to assess effects of low dose, acute Wnt inhibition. **c**. Images of cells treated with 0.4uM CHIR99021 as indicated in **b**, after 9 days of differentiation. Scale bars, 50 µm. **d**. Lipid droplet quantification of cells illustrated in **c** (error bars = SD, *n =* 3, \**p*-value ≤ 0.05, determined by one-way ANOVA). **e**,**f**. *ADIPOQ* and *MGP* mRNA levels of cells treated with 0.4uM CHIR99021 as indicated in **b**, after 3 days of differentiation (error bars = SD, *n* = 2, exact p-values shown were determined by one-way ANOVA). **g**. Images of cells treated with 0.4uM XAV939 as indicated in **b**, after 9 days of differentiation. Scale bars, 50 µm. **h**. Lipid droplet quantification of cells illustrated in **g** (error bars = SD, *n =* 3, exact p-values shown were determined by one-way ANOVA). **i**,**j**. *ADIPOQ* and MGP mRNA levels of cells treated with 0.4uM XAV939 treatment measured by qRT-PCR after 3 days of differentiation (error bars = SD, *n* = 2, exact p-values shown were determined by one-way ANOVA).

## DISCUSSION

Multipotent mesenchymal progenitor cells are required throughout lifetime to renew and repair multiple tissues. However, how these progenitor cells are maintained under differentiation signals is not known. A striking finding in our study is that primary mesenchymal progenitors derived from human adipose tissue are highly homogenous at single-cell resolution. However, after only 3 days of adipogenic induction, clear heterogeneity develops and the cells diverge towards two trajectories; one toward the adipocyte phenotype, and another toward an undifferentiated state that maintains proliferative capacity and multipotency. These findings reveal that induction of adipogenic differentiation is coupled to a mechanism to preserve a reservoir of multipotent progenitor cells, allowing for tissue maintenance and development throughout the lifetime.

The discovery of this mechanism required a model to examine the earliest stages of human adipocyte development at scale, as the turnover rate of adipocytes in adult humans is exceedingly low, and therefore developmental trajectories cannot be captured at steady-state ^7^. Indeed, in single-cell/single-nuclei profiles of human adipose tissues, progenitors and mature adipocytes are projected as distinct spatial clusters but the developmental trajectory of adipocytes is not well captured ^29,30^,

A second key finding in our study is that the generation of cells in the reservoir pool is regulated expression of canonical Wnt target genes, including the specific marker *MGP*. Transcriptomic profiles of early multi-lineage differentiated cells revealed the *MGP*^+^ cells also developed in response to osteogenic and chondrogenic induction (Extended Data Fig. 3d), implying the mechanism that actively maintains a multipotent mesenchymal progenitor cell pool is not unique to the adipose lineage. Through meta-analysis of internally generated and published transcriptomic data, we provide evidence that *MGP*^+^ cells are present in ASPCs in human adult adipose tissue and they are likely maintained throughout lifetime. Moreover, in agreement with our observations, a recently posted ^31^ single-cell profiling study on adipose progenitors harvested from four different human adipose tissue depots find that these cells undergo two trajectories upon adipogenic induction. Thus, *MGP*^+^ mesenchymal progenitors with ability to proliferate and retain multipotent potential are likely to be found in diverse human depots.

Our study identified canonical Wnt signaling as a key factor for maintaining cells in a multipotent progenitor state. Much of our understanding of Wnt signaling comes from its role in epithelial cells, particularly in the intestine, and on the effects of aberrant Wnt signaling on the development of cancer ^32,33^. In humans, nineteen *Wnt* ligands, belonging to twelve subfamilies, may act upon Wnt receptor complex composed of ten possible *Frizzled* proteins and *LRP5/6* receptors. Wnt-driven gene expression mediated by the *β-Catenin/TCF* transcription factors is collectively described as canonical Wnt signaling, and is dependent on intracellular regulators including *Dishevelled, Axin, GSK3* and *APC*, as well as extracellular regulators including *Dickkopf*-family (*DKK*) and *Secreted Frizzled Related Protein*-family (*SFRP*) proteins.

The Wnt signaling pathway elements involved in maintaining a pool of multipotent cells are inherently expressed in response to adipogenic stimuli, as no exogenous Wnt perturbagens were applied in our study. These signaling mechanism operate very early in the process of determination, as brief and mild perturbation of canonical Wnt activity early in adipogenic induction was sufficient to alter fate preference between progenitor maintenance and differentiation. Our results are consistent with the important role of Wnt in stem/progenitor cell maintenance in multiple tissues and organs ^1-3^. Existing studies of the role of Wnt in adipogenesis, first described in 2000 ^12^, were extensively reviewed recently by de Winter and Nusse ^34^, who note that inactivation of Wnt signaling is necessary for mesenchymal progenitor lineage commitment, and further postulate that Wnt inhibits adipogenesis by promoting mesenchymal progenitor maintenance. Our findings provide direct evidence for that hypothesis, showing that Wnt signaling preserves cells in a progenitor state even in the context of strong adipogenic induction.

Potential Wnt-dependent interactions between cells undergoing the adipocyte versus multipotent trajectories upon stimulation can be hypothesized based on the expressed ligand-receptor pairs. One of these is the *Ephrin B/Ephrin B receptor* pathway, where *Ephrin B* from *ADIPOQ*^+^ adipocytes can signal to *Ephin B receptor*-expressing *MGP*^+^ cells. Wnt-mediated *Ephrin* signaling is known for directing development and cell positioning in the intestinal epithelium ^1^. The spatial relationship between adipocytes and mesenchymal progenitors’ niche are challenging to model, but Merrick at al. ^27^ find that in mice, progenitors expressing the Wnt targe gene *Dpp4* reside in a specific, extracellular matrix-rich region of adipose tissue, from where they are mobilized under conditions of tissue remodeling. Mouse *Dpp4+* progenitor cells might be analogous to human *MGP+* cells, and reflect a reservoir of multipotent progenitors formed during tissue development that serve to repair and replenish the tissue during the life of the mouse. Other Wnt genes identified are secreted factors and regulators of Wnt signaling, including *SFRP2* and *DKK1*, which may serve as a paracrine signal promoting adipogenesis and an autocrine negative feedback signal preventing overactivation of Wnt signaling. Further studies will focus on developing models for analyzing the topology of adipose tissue development and the underlying autocrine mechanisms.

An additional observation made during our experiments is that Wnt has a potentially different role in supporting adipocyte development post-fate commitment. While brief inhibition of *Tankyrase* during adipogenic induction promotes adipogenesis, chronic inhibition of *Tankyrase*, which stabilizes *Axin* and suppresses *β-catenin* activity ^35^, strongly suppresses adipocyte development without inducing toxicity (Extended Data Fig. 7). We hypothesize Wnt is also important for adipocyte development after fate determination, which is supported by the report of *β-catenin* requirement in adipose tissue lipogenesis demonstrated in an adipocyte-specific *β-catenin* knockout mouse model ^10^. The distinct mechanisms by which Wnt signaling controls fate determination early upon differentiation induction and supports adipocyte development post-fate commitment remain to be elucidated.

Our findings reveal orchestrated Wnt signaling at different stages of adipose tissue development. Variations in Wnt activity may be critical in determining mechanisms of adult adipose tissue expansion. As Wnt signaling is driven by specific combinations of Wnt ligand, extracellular and intracellular regulators, and receptor complexes, further characterization of adipose-context specific Wnt signaling may reveal opportunity for developing therapeutic interventions for improving adipose tissue function.

## MATERIALS & METHODS

### Adipose tissue explants

Methods for the harvesting and the culture adipose tissue explants were previously published (Min, et al., 2016). In short, subcutaneous adipose tissues were donated from consented adult patients (demographics listed in table below) undergoing elective panniculectomy surgery at University of Massachusetts Medical Center under UMass Institutional Review Protocol #14734_13 and were subjected to harvesting within one to six hours. 200 explants of approximately 1cm^3^ in size were embedded in Matrigel Matrix (Cat# 356231, Corning) per 10 cm dish with EGM-2MV (Cat# CC-3156,CC-4147, Lonza) media supplementation. The progenitors in explants were allowed to grow for 14 days in Matrigel with fresh media replacement every 2-3 days. After 14 days, progenitors in explants were recovered using Dispase (Cat# 354235, Corning) for one hour followed by additional 14 minutes of Trypsin-EDTA (Cat# 15400-054, Gibco) and Collagenase I (Cat# LS004197, Worthington) and plated on standard tissue culture plate for expansion and cryopreservation.

| Donor # | Age | Gender | BMI |
| --- | --- | --- | --- |
| 37 | 57 | Female | 37.8 |
| 45 | 41 | Female | 32.9 |
| 48 | 59 | Female | 27.6 |
| 49 | 47 | Female | 24.3 |
| 57 | 41 | Female | 37.8 |
| 62 | 60 | Female | 38.3 |
| 65 | n/a | n/a | n/a |
| 68 | 33 | Female | 28.6 |

### Lineage differentiation

Adipogenic differentiation was induced by providing confluent cells with DMEM (Cat# 11995-065, Gibco) and 10% Fetal Bovine Serum (FBS)(Cat# 25-514, Genesee Scientific) supplemented with 0.5 mM 3-isobutyl-1-methylxanthine (Cat# I5879, Sigma), 0.25 μM dexamethasone (Cat# D1756, Sigma), and 5 μg/mL insulin (Cat# 15500, Sigma). The induced cells underwent half media replacement every 24 hours for 3 days. After 3 days, the media is replaced with fresh DMEM + 10% FBS media every 2-3 days until harvest.

Chondrogenic differentiation was induced by providing confluent cells with DMEM and 10% FBS supplemented with 1 mM sodium pyruvate (Cat# 11360-070, Gibco), 100 nM dexamethasone (Cat# D1756, Sigma), 10 ng/mL Human TGF**-**β 1 recombinant protein (Cat# PHG9204, Gibco) and 1 μg/mL L-Ascorbic acid 2-phosphate (Cat# A8960, Sigma). The induced cells underwent half media replacement every 2-3 days until harvest.

Osteogenic differentiation was induced by providing confluent cells with DMEM and 10% FBS supplemented with 10 mM sodium beta-glycerolphosphate (Cat# L03425, Alfa Aesar), 100 nM dexamethasone (Cat# D1756, Sigma), 50 uM L-Ascorbic acid 2-phosphate (Cat# A8960, Sigma). The induced cells underwent half media replacement every 2-3 days until harvest.

Cells in the control condition were maintained in DMEM + 10% FBS and underwent identical media replacement schedule as the experimental group.

### Lineage staining

Cells were washed with PBS and fixed with 10% formalin for 30 minutes at room temperature. Following fixation, cells were washed three times with double distilled water. To stain adipocyte lipid droplets, cells were first incubated in 60% isopropanol for 5 minutes followed by staining with 2% Oil Red O (Cat# 00625, Sigma) in 60% isopropanol for 10 minutes. For assessment of chondrogenesis, cells were incubated with 1% Alcian Blue 8GX (Cat# A5268, Sigma) in 2:3 acetic acid and ethanol solution in the dark for 30 minutes with gentle agitation. For assessment of osteogenesis, cells were incubated with 2% Alizarin Red S staining solution (Cat# 0223, ScienCell) in the dark for 30 minutes with gentle agitation. After each staining protocol, the staining solution was removed, and cells were washed three times with double distilled water before imaging.

### RNA extraction for bulk RNA-sequencing and qPCR

Cells in culture wells were washed with PBS before harvesting with TriPure TRIzol reagent (Cat# 11 667 165 001, Roche). The cell-TRIzol mixtures were transferred to collection tubes and homogenized with Tissuelyser II (Qiagen). Chloroform was added in a 1:5 ratio by volume and phase separation was performed. The RNA-containing layer was mixed with an equal volume of 100% isopropanol and incubated overnight at -20 °C for precipitation. RNA was pelleted and washed with 80% ethanol and resuspended in nuclease-free water. RNA concentration and purity were determined using a NanoDrop 2000 (Thermo Scientific). RNA for sequencing were sent to University of Massachusetts Medical School Molecular Biology Core Lab for fragment analysis.

### Bulk RNA-sequencing

Library preparation was performed using TruSeq Stranded mRNA Low-Throughput Sample Prep Kit (Cat# 20020594, Illumina) according to manufacturer’s instruction. The libraries were sequenced on the NextSeq 500 system (Illumina) using the NextSeq® 500/550 High Output Kits v2 (75 cycles; single-end sequencing; Cat# FC-404-2005, Illumina). The FASTQ files were processed using the DolphinNext pipeline ^36^ on the Massachusetts Green High Performance Computer Cluster (GHPCC). DolphinNext was configured to use RSEM for read mapping and transcript quantification ^37^. Differentially expressed genes were identified using DESeq2 ^38^. Pathway analysis was performed using the Gene Set Enrichment Analysis (GSEA) software with the MSigDB GO biological process gene sets (http://www.gsea-msigdb.org/gsea/msigdb/annotate.jsp) ^39^. Sequencing results were submitted to GEO (accession number: GSE198275, GSE198481, GSE204847, GSE204848) and will be made publicly available upon publication of the manuscript.

### Single cell RNA-sequencing

Single-cell library preparation was performed using Chromium™ Single Cell 3’ GEM Library & Gel Bead Kit v3 (Cat# 1000092, 10X Genomics) according to manufacturer’s instruction. The libraries were sequenced on the NextSeq 500 system (Illumina) using the NextSeq® 500/550 High Output Kits v2.5 (50 cycles; Cat# FC-404-2005, Illumina). The sequencing outputs were processed using the CellRanger software v3.1.0 on the Massachusetts Green High Performance Computer Cluster (GHPCC), Reads were mapped to human reference genome GRCh38 (Ensembl 93). Data analysis was performed using Seurat v4.1.0 ^40^ within R version 4.0.2 environment. RNA velocity analysis was performed using the velocyto v0.17 commend line tool and velocyto.R v0.6 R package ^41^. Sequencing results were submitted to GEO (accession number: GSE198482). Sequencing results and analysis scripts will be made publicly available upon publication of the manuscript.

### Quantitative PCR with reverse transcription (qRT-PCR)

1 µg of RNA was reverse transcribed using the iScript cDNA Synthesis Kit (Cat# 1708891, Bio-Rad) according to manufacturer’s protocol. Quantitative reverse-transcription PCRs were prepared with iQTM SYBR Green Supermix (Cat# 1708882, Bio-Rad) and were performed on a CFX Connect Real-Time PCR Detection System (Bio-Rad). The ADIPOQ primers have the following sequences: 5’-TGC TGG GAG CTG TTC TAC TG-‘3 forward and 5’-TAC TCC GGT TTC ACC GAT GTC-‘3 reverse. The MGP primers have the following sequences: 5’- CAG CAG AGA TGG AGA GCT AAA G -‘3 forward and 5’- GTC ATC ACA GGC TTC CCT ATT -‘3 reverse. The FRZB primers have the following sequences: 5’- GCC CTG GAA CAT GAC TAA GAT G -3’ forward and 5’- GTA CAT GGC ACA GAG GAA GAA G -’3 reverse. The COMP primers have the following sequenes 5’- CCA ACT CAA GGC TGT GAA GTC -‘3 forward and 5’- GGA CTT CTT GTC CTT CCA ACC -3’ reverse.

### Image acquisition and processing

Cells were imaged with LEICA DM 2500 LED inverted microscope equipped with a Leica MC120 HD digital camera. Fiji/ImageJ v1.53c software was used to quantify lipid droplets. The images were converted from RGB to 8-bit, background subtracted, contrast enhanced, thresholded and binarized followed by circular particle analysis (Extended Data Fig. 8).

### Cell separation with Percoll density gradient

A Percoll step density gradient was prepared in a 15ml conical tube with the Percoll solutions (Cat# P4937, Sigma). Earlier experiments (Fig. 4a-c) were performed in Percoll densities of 1.010 g/mL, 1.020 g/mL and 1.040 g/mL, while later experiments were performed in Percoll densities of 1.010 g/mL, 1.020 g/mL and 1.030 g/mL to permit high density cells to be collected from the cell pellet after centrifugation.

7-day adipogenic induced cell populations were lifted with StemPro Accutase (Cat# A1110501, ThermoFisher). Lifted cells were pelleted and resuspended in 1.010 g/mL Percoll solution and loaded onto the top of the prepared Percoll Gradient, followed by centrifugation at 1000g for 30 minutes at room temperature. Cell fractions were observed by eye and each resulting fraction was pipetted into new conical tubes for further experimentations.

### Ligand-receptor analysis

First, we identified genes expressed in the *MGP*^*+*^ and *ADIPOQ*^*+*^ clusters. A gene was considered as expressed within a cell cluster if average normalized counts >= 0.5. We then queried the putative or literature supported ligand-receptor pairs obtained from Ramilowski, et al., 2015 to identify gene pairs expressed in the *MGP*^*+*^ and *ADIPOQ*^*+*^ clusters.

### Cell viability assays

CellTiter-Glo 2.0 cell viability assay (Cat# G9243, Promega) and CellTox Green cytotoxicity assay (Cat# G8742, Promega) were performed according to manufacturer’s instruction and the fluorescence and luminescence signals were measured using a Safire 2 microplate reader (Tecan).

### ELISA

Adiponectin concentration in the conditioned medium was measured using the Adiponectin Human ELISA Kit (Cat# KHP0041, ThermoFisher) with a Safire 2 microplate reader (Tecan).

### Small molecule inhibitors

Tankyrase inhibitor XAV-939 (Cat# HY-15147), GSK3 inhibitor CHIR-99021 (Cat# HY-10182), and DPP4 inhibitors MK-043 (Cat# HY-13749) and LAF237 (Cat# HY-14291) were obtained from MedChemExpress.

## CODE AVAILABILITY STATEMENT

Code used in this study for the analysis of the single-cell RNA-seq data will be made available at https://github.com/zingery/mesenchymal-maintenance/.

## ACKNOWLEDGEMENTS

We thank MedChemExpress for their generosity in providing the Wnt compounds XAV939 and CHIR99021. This study was supported by NIH grants DK089101 and DK123028 to SC and GM135751 to JSR. We acknowledge the UMass Chan IT department for computing infrastructure.

## AUTHOR CONTRIBUTIONS

ZYL: conceptual design, experiment design and performance, data analysis, manuscript preparation. SJ: conceptual design and performance. JSR: hypothesis generation, conceptual design, experiment design and performance, data analysis, manuscript preparation. AD: experiment design and performance, data analysis. PS: conceptual design and performance. QY: conceptual design, manuscript preparation. TD: experiment design and performance, data analysis, manuscript preparation. TN: conceptual design, manuscript preparation. OAM: conceptual design, data generation, manuscript preparation. SC: supervision of work, hypothesis generation, conceptual design, manuscript preparation.

## EXTENDED DATA FIGURES

**Extended Data Fig. 1.**
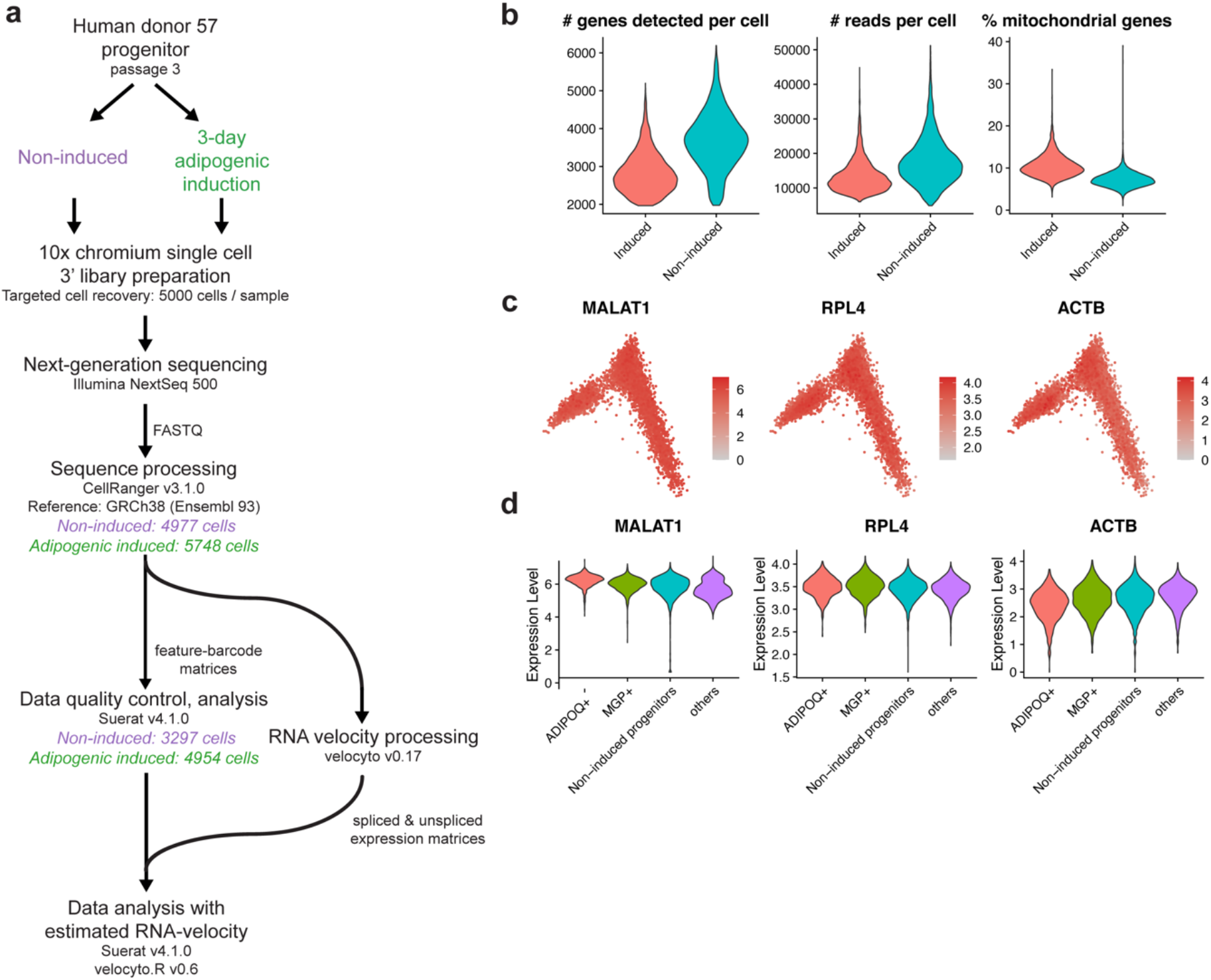
Processing and quality control of the single-cell RNA-seq profiling of non-induced and 3-day adipogenic induced progenitors. **a**. Schematic of the sample and bioinformatics processing of the study. **b**. Violin plots of the number of genes/reads detected and estimated percentage of mitochondrial genes in cells that had passed quality control. **c** Selected housekeeping genes’ expression in PCA projection. **d**. Violin plot of gene expression distribution of selected housekeeping genes.

**Extended Data Fig. 2.**
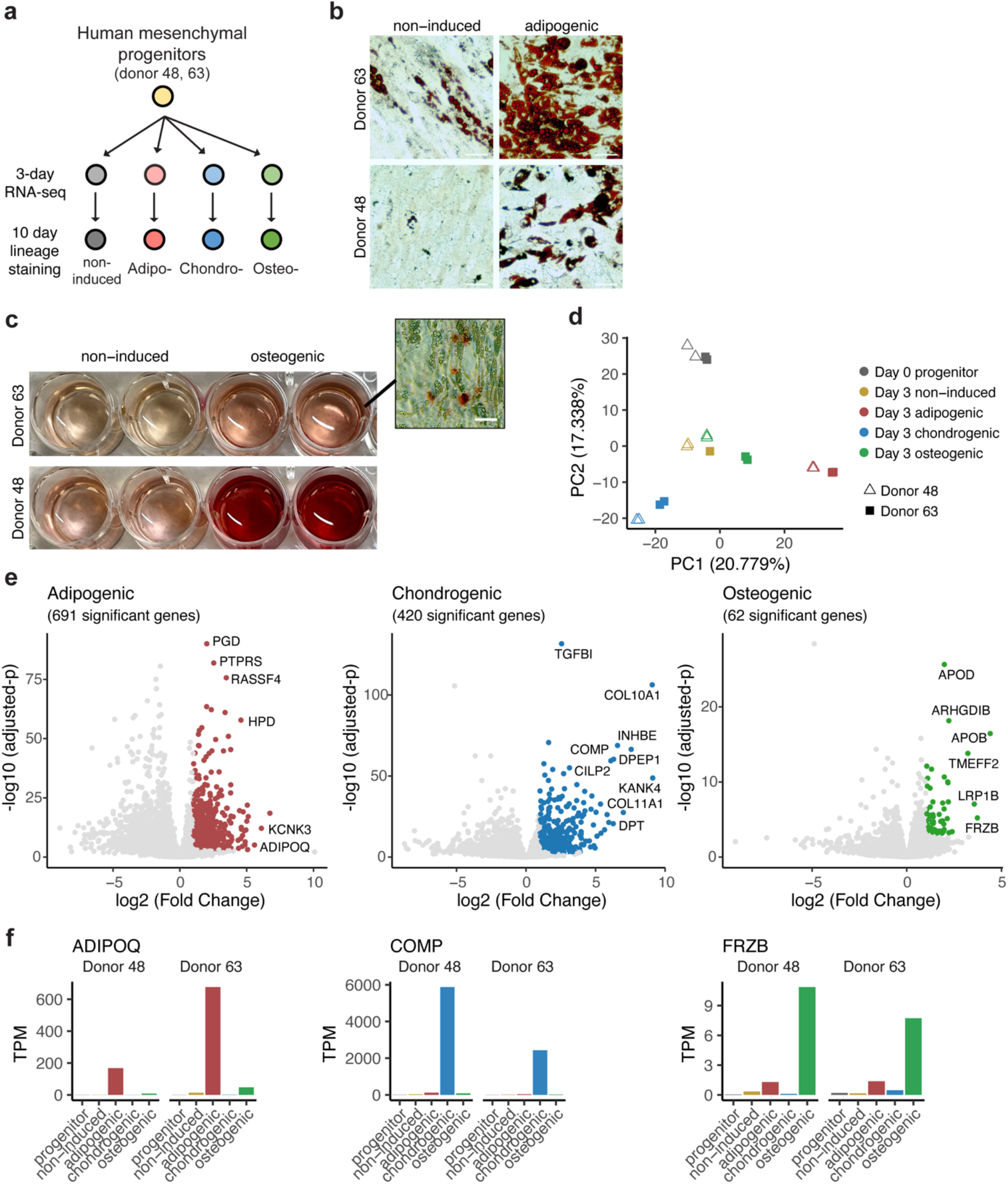
Identification of early osteogenic and chondrogenic lineage marker with multi-lineage time course bulk RNA-seq. **a**, Schematic of the RNA-seq profiling study. Two donor-derived cells separately expanded at two biological replicates were used to obtain RNAs at 3 days post differentiation induction for bulk RNA-sequencing. **b**, Oil Red O staining of progenitor cells underwent 10-day adipogenic induction. Scale bars, 50 µm. **c**, Alizarin Red S staining of progenitor cells underwent 10-day osteogenic induction. **d**, Scatter plot of the first two principal components of the RNA-seq samples. Principal component analysis was performed on the expression of the top 1000 most variable genes across all samples. **e**, Volcano plots of differential gene expression analysis results of the transcriptome profiles between cells induced toward the annotated lineage and other 3-day induced samples. Differentially expressed genes were defined as those with log2 fold change > 1 and adjusted p-value < 0.001. **f**, Gene expression profile of distinctive lineage marker identified from the differential expression analysis. Markers were selected based on magnitude and significance in the differential expression analysis as well as specificity. *ADIPOQ: Adiponectin, COMP: Cartilage Oligomeric Matrix Protein, FRZB: Frizzled Related Protein*.

**Extended Data Fig. 3.**
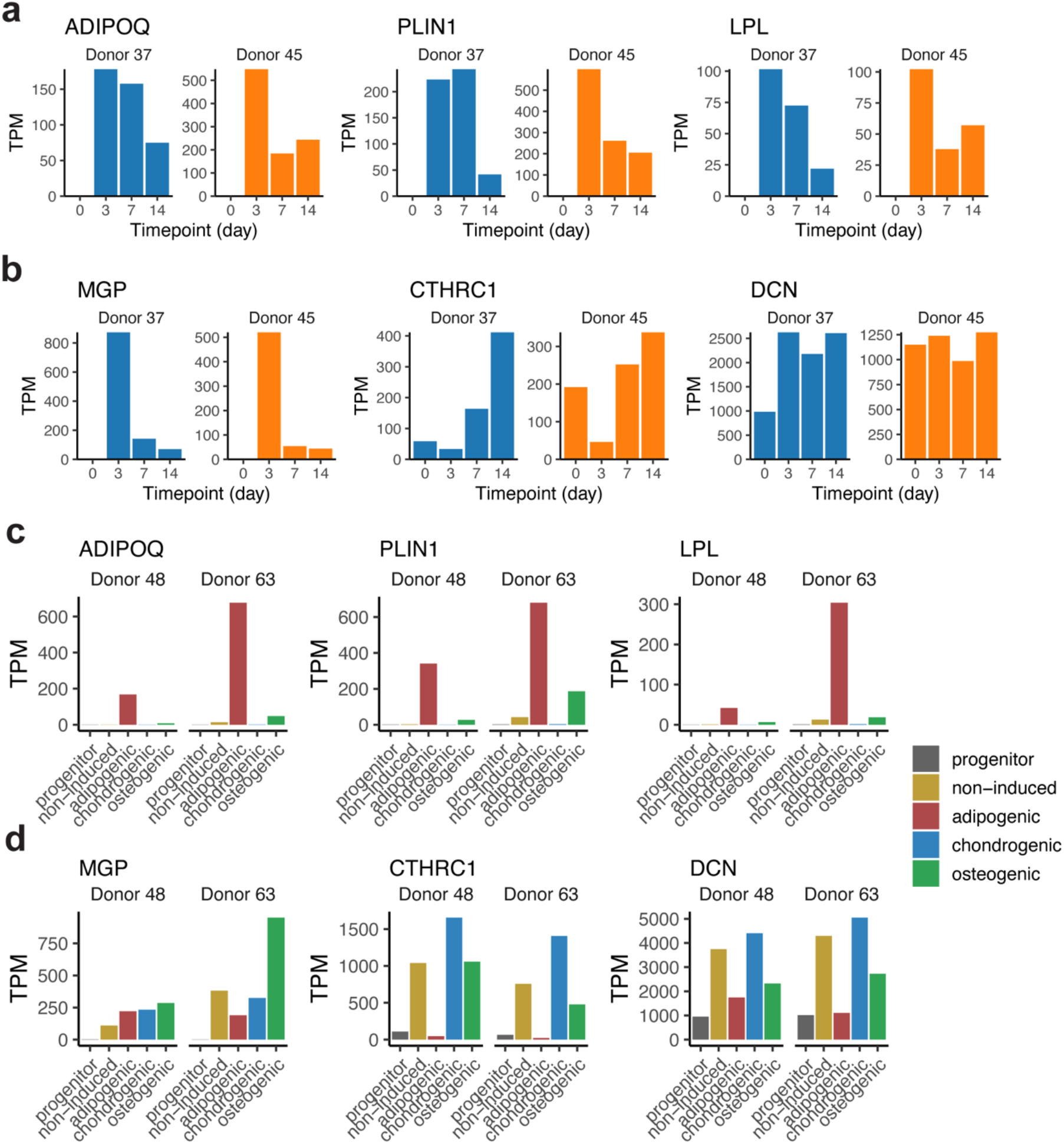
Gene expression profiles of markers identified from the single-cell RNA-seq or multi-lineage bulk RNA-seq datasets of induced adipose progenitors in adipogenesis time course. **a**. Gene expression profiles of *ADIPOQ*^+^ cell markers in the adipogenesis time course RNA-seq dataset presented in Fig. 1d. **b**. Gene expression profiles of *MGP*^+^ cell markers in the adipogenesis time course RNA-seq dataset presented in Fig. 1d. **c**. Gene expression profiles of *ADIPOQ*^+^ cell markers in the 3-day multi-lineage RNA-seq dataset described in Extended Data Fig. 2. **d**. Gene expression profiles of *MGP*^+^ cell markers in the adipogenesis time course RNA-seq dataset described in Extended Data Fig. 2.

**Extended Data Fig. 4.**
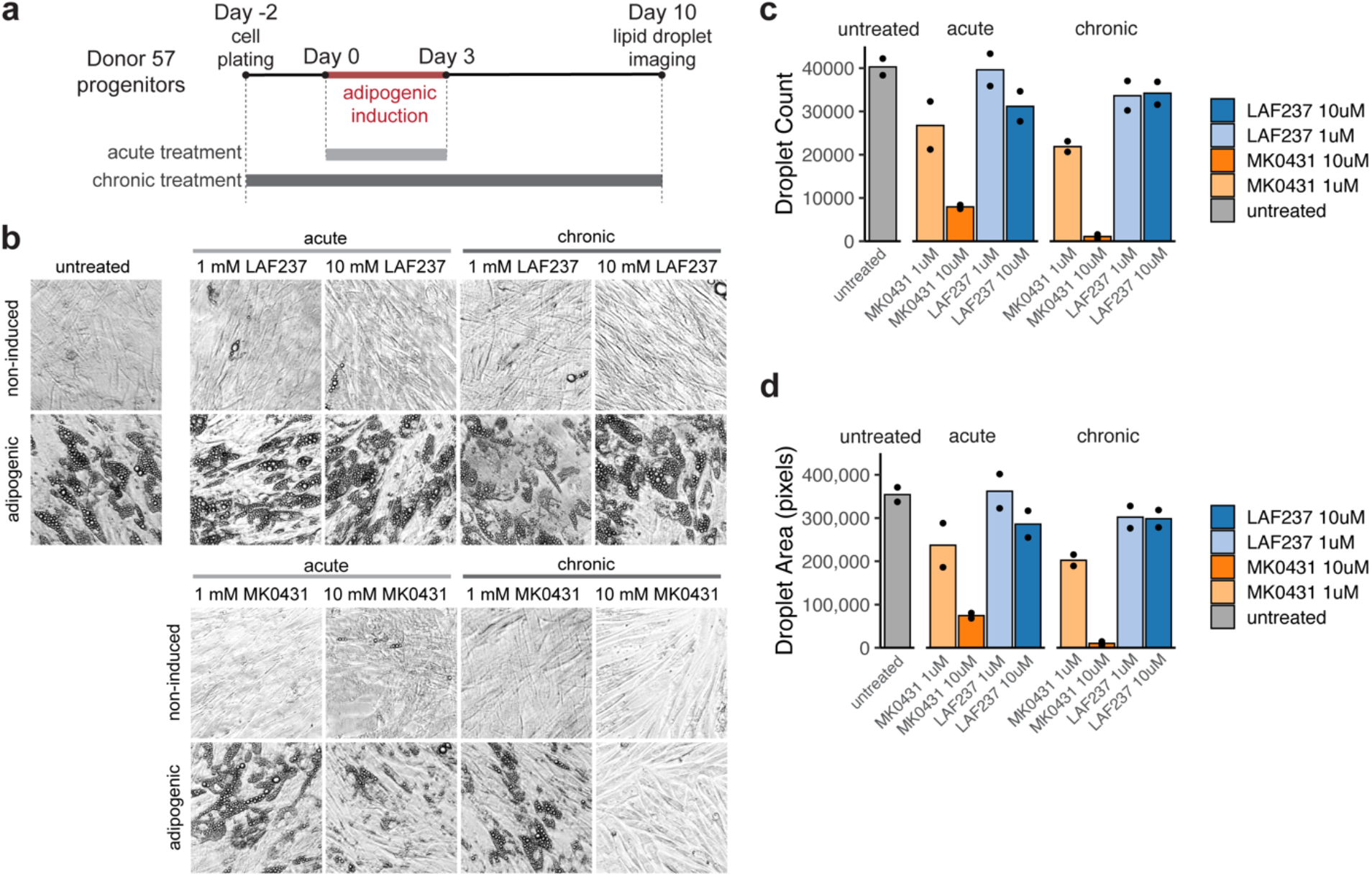
*DPP4* inhibition does not promote adipogenesis. **a**. Schematic of the adipogenesis assay with acute or chronic *DPP4* inhibitor treatment, n=2. **b**, Images of 10-day adipogenic induced cells with acute or chronic *DPP4* inhibitor treatment. **c**,**d**, Lipid droplet quantification of 10-day adipogenic-induced cells with acute or chronic *DPP4* inhibitor treatment.

**Extended Data Fig. 5.**
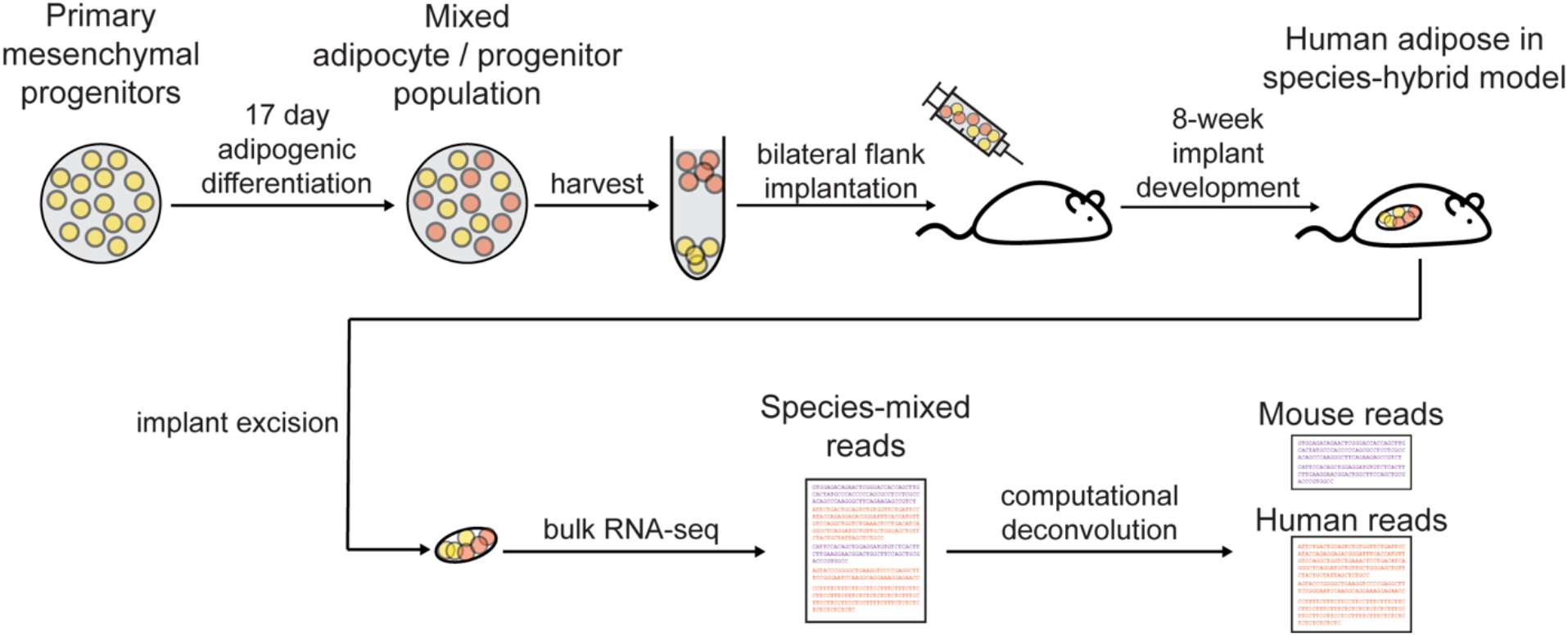
Human adipogenic-induced progenitor mouse implantation model.

**Extended Data Fig. 6.**
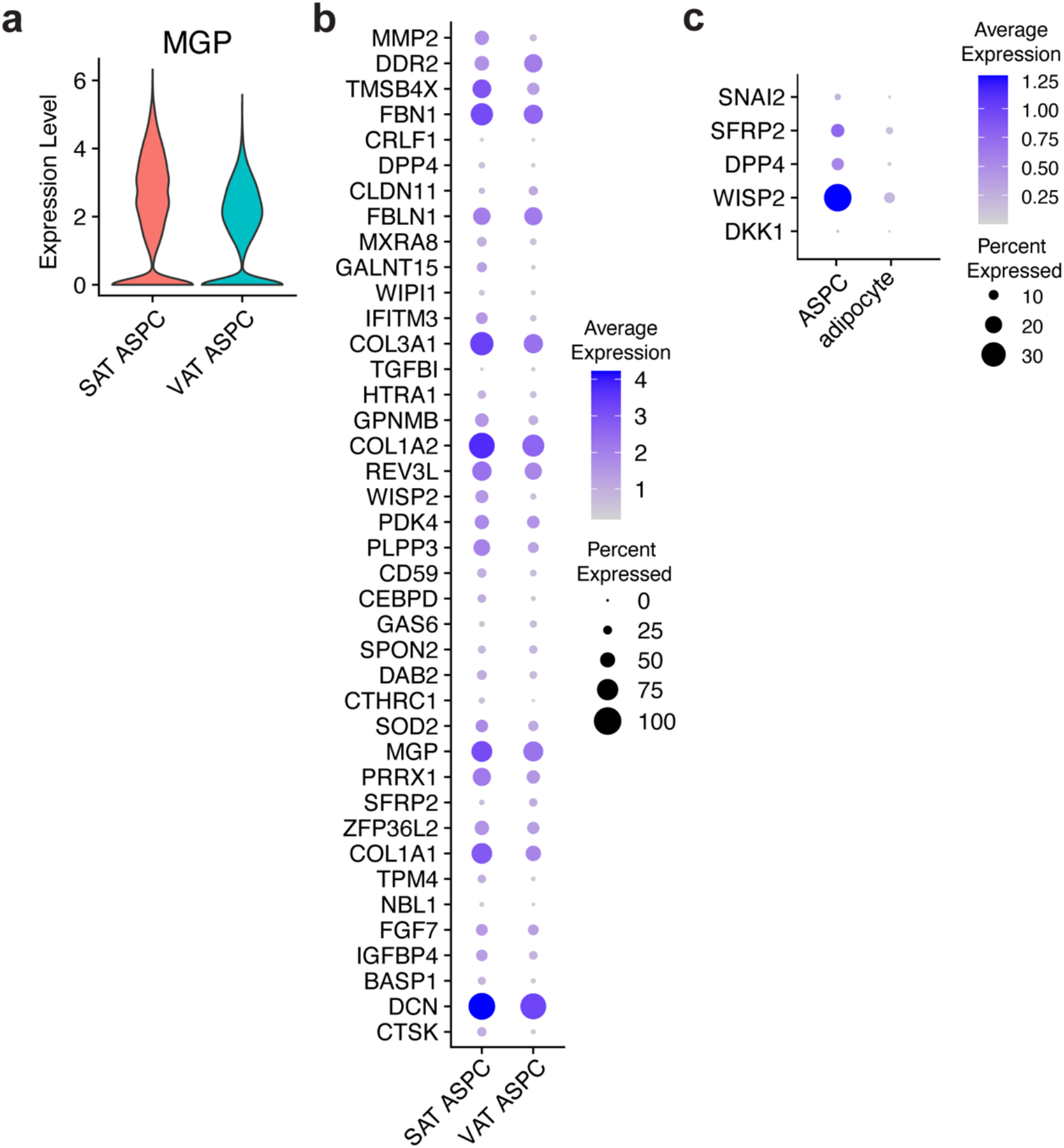
*MGP*^+^ cells resemble human adult APSCs and are enriched in subcutaneous adipose tissue comparing to visceral adipose tissue. **a**. Violin plot of *MGP* expression visualized by adipose tissue depot from the Emont, et al. dataset. **b**. Dot plot of the top 40 *MGP*^***+***^ markers gene expression visualized by adipose tissue depot from the Emont, et al. dataset. **c**. Canonical Wnt target genes identified in *MGP*^+^ cells queried from the Emont, et al. dataset.

**Extend Data Fig. 7.**
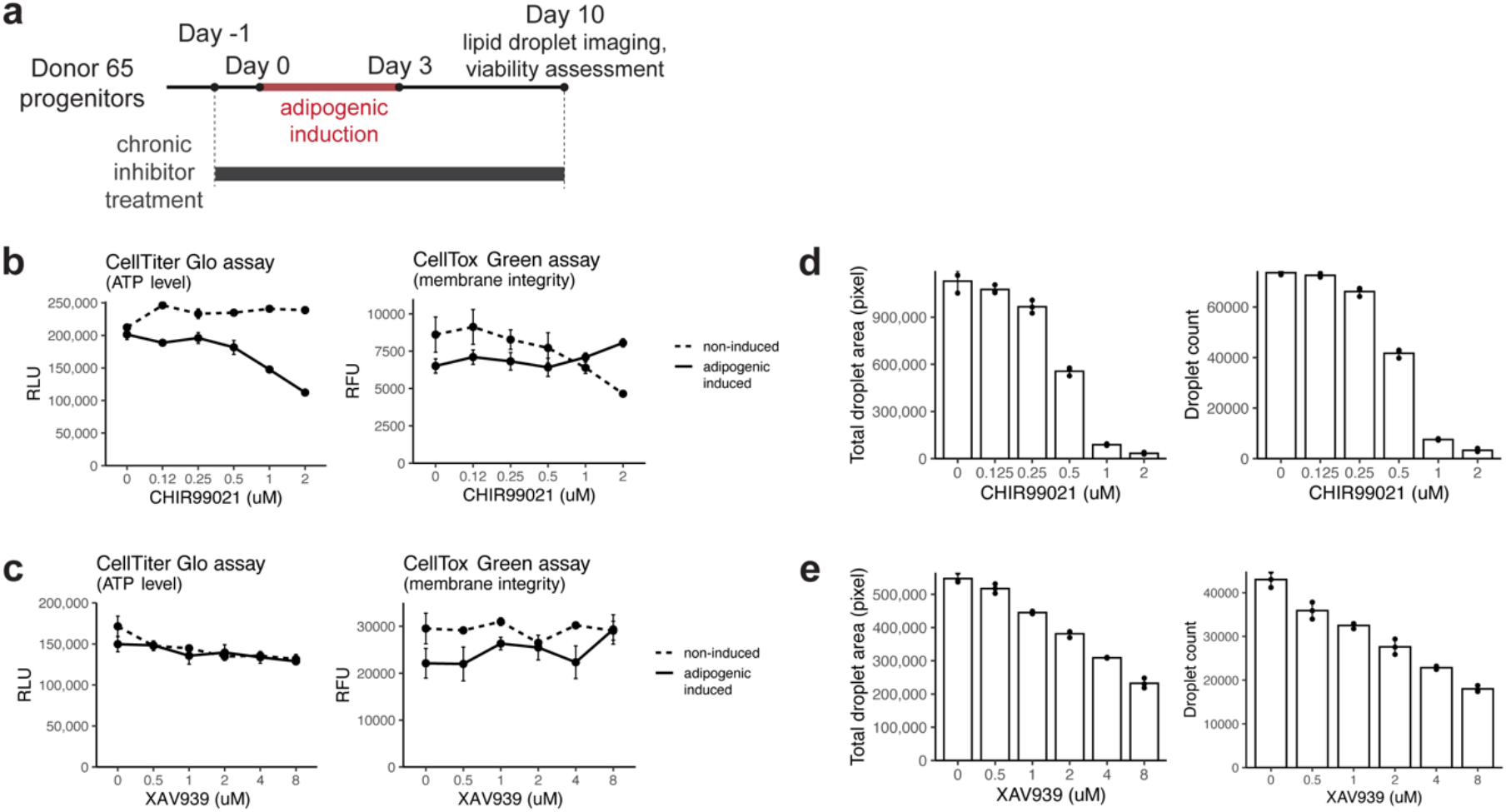
Long term Wnt agonist/antagonist treatment suppresses adipogenesis. **a**, Schematic of the adipogenesis assay with chronic Wnt inhibition. **b**. Viability assessment of 10-day adipogenic-induced cells under chronic exposure to different dosages of CHIR99021 (error bars = SD, *n =* 3). **c**. Viability assessment of 10-day adipogenic-induced cells under chronic exposure to different dosages of XAV939 (error bars = SD, *n =* 3). **d**. Lipid droplet quantification of 10-day adipogenic-induced cells under chronic exposure to different dosages of CHIR99021 (error bars = SD, *n =* 3). **e**. Lipid droplet quantification of 10-day adipogenic-induced cells under chronic exposure to different dosages of XAV939 (error bars = SD, *n =* 3).

**Extended Data Fig. 8.**
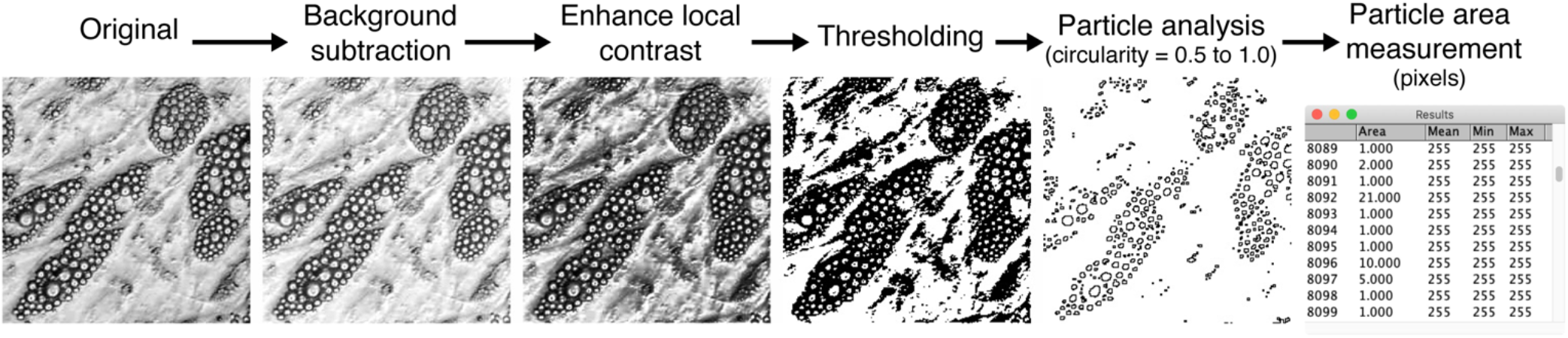
Lipid droplet quantification permits statistical comparison of lipid number and size. **a**. Schematic of image processing process for lipid droplet quantification from microscopy images.

